# Pair-wise Comparison Analysis for Multiple Pool-seq: an efficient method identified anthocyanin biosynthesis genes in rice pericarp

**DOI:** 10.1101/561258

**Authors:** Xinghai Yang, Xiuzhong Xia, Zongqiong Zhang, Baoxuan Nong, Yu Zeng, Yanyan Wu, Faqian Xiong, Yuexiong Zhang, Haifu Liang, Yinghua Pan, Gaoxing Dai, Guofu Deng, Danting Li

**Affiliations:** Rice Research Institute, Guangxi Academy of Agricultural Sciences, Nanning, China; Biotechnology Research Institute, Guangxi Academy of Agricultural Sciences, Nanning, China; Cash Crops Research Institute, Guangxi Academy of Agricultural Sciences, Nanning, China

**Keywords:** multiple pools, whole genome resequencing, SNP-index, anthocyanin, candidate genes.

## Abstract

The complex traits are derived from multiple genes and exhibit a large variety of phenotypes. High-throughput sequencing technologies have become the new strategies for mapping the important traits of crops. However, these methods have their own disadvantages and limitations. Here we introduced Pair-wise Comparison Analysis for Multiple Pool-seq (PCAMP) for mapping the candidate genomic regions involved in anthocyanin biosynthesis in rice pericarp. In this protocol, the second filial generation (F_2_) populations obtained by crossing two parents with different target traits were divided into n (n>=3) subpopulations according to their phenotypes. Thirty phenotypically identical individuals were selected from each subpopulation and DNA samples were extracted to form a pool for sequencing. Finally, we compared the SNP-index between every two Pool-seqs to map the candidate genomic regions. We applied PCAMP to analyse F_2_ populations and successfully identified five known genes and five new candidate genomic regions for anthocyanin biosynthesis in rice pericarp. These results demonstrate that PCAMP is an efficient new method for dissecting the complex traits of crops.

---

The world population is likely to grow by 60%–80% within the next 50 years, which requires the food production to nearly double, based on current living standards. Due to global warming and the increasing reduction of arable land, increasing grain production is a great challenge(Abe et al., 2012). The complex traits of crops are important for adaption to environment and genetic improvement(Mitchell-olds, 2010), which are genetically controlled by multiple genes. Each gene has relatively minor effect on phenotype and is susceptible to environmental influences(Buckler et al., 2009). Therefore, it is quite difficult to dissect the genetic basis underlying the complex traits.

BSA (Bulked Segregant Analysis) is a approach for map inherited traits. Initially, BSA was used to identify molecular markers closely linked to target genes in lettuce(Michelmore et al., 1991) and tomato(Giovannoni et al., 1991). Subsequently, BSA combined with RFLP, RAPD, SSR and other molecular markers was used to reveal target genes in soybean(Darvasi a, 1994; Mansur et al., 1993). With the development of high throughout sequencing technology, Schneeberger et al. (Singh et al., 2017) identified the candidate gene AT4G35090 which leads to slow growth and light green leaves of Arabidopsis thaliana by SHOREmap technology. Austin et al. (Austin et al., 2011) invented a BSA method based on homozygosity mapping, which is mainly applied to map recessive traits. Ehrenreich et al.(Ehrenreich et al., 2010) and Wenger et al. (Wenger et al., 2010)successfully located target genes in yeast and Saccharomyces cerevisiae respectively. Recently, Abe et al. (Abe et al., 2012)used MutMap technology to locate rice green leaf mutant gene *OsCAO1*. Fekin et al.(Fekih et al., 2013) developed MutMap+ technology, which was applicable to early mutagenesis and lethal or non-inbred mutants. based on MutMap and de novo assembly, Takagi et al.(Takagi et al., 2013A)developed a new mapping approach, MutMap-Gap applicable to the mutation sites which were not found in the reference genome. In 2013, Takagi et al. (Takagi et al., 2013B) first used QTL-seq technology to locate rice blast and seedling vigor related genes. Since then, the BSA has become a common approach for locating genes.

With the application of QTL-seq technology in plant trait mapping, many new methods have been developed. Das et al.(Das et al., 2016) crossed a less pod number variety ILWC 46 with two more pod number varieties Pusa 1103 and Pusa 256 respectively to obtain the F2 population for mQTL-seq, and then narrowed the candidate region through taking the intersection of the two candidate intervals. Kumar et al.(Kumar et al., 2018) identified three genes related to Ascochyta blight resistance of chickpea based on mQTL-seq technology. The application of QTL-seq to some large genome, such as barley, is challenging, so Hisano et al.(Hisano et al., 2017)identified the monogenic Mendelian locus and associated QTLs using Exome QTL-seq. Xue et al.(Xue et al., 2017) modified the QTL-seq method by introducing a | ((SNP-index)| parameter to improve the accuracy of mapping the red skin trait in a group of highly heterozygous Asian pears. Yoshitsu et al.(Yoshitsu et al., 2017) constructed Y-bulk (early-heading), S-bulk (late-heading) and L-bulk (extremely late-heading) from F2 population of two foxtail millet cultivars lines, and then identified two QTLs associated with DTH by genome-wide comparison of SNPs in the Y-bulk versus the S-bulk and the Y-bulk versus the L-bulk. Huang et al.(Huang et al., 2017)identified a Clubroot resistance related gene in Chinese Cabbage using BSR-Seq technology. Dou et al.(Dou et al., 2018) obtained a watermelon shape related candidate region through GWAS analysis, and further validated and narrowed candidate interval based on BSA-seq and genetic linkage analysis. Singh et al.(Singh et al., 2017) constructed two mixed pools using the F2 population of ICPL 20096 (resistant to fusarium wilt and sterility mosaic disease) and ICP 332 (susceptible to FW and SMD), and located FW and SMD resistance related genes through indel-seq approach.

QTL-seq and other derivatives play an important role in plant trait mapping. However, these methods only select plants with extreme phenotypes of target traits in the segregated population and identify one or two genes/QTLs associated with related traits. In crops, many agronomic traits are complex traits controlled by multiple genes. The traditional method of quantitative trait analysis is only for single QTL and time-consuming and labor intensive cost(Tuberosa et al., 2005). GWAS analysis can be used to quickly identify complex traits related candidate intervals, but it has limitations(Platt et al., 2010; Farlow, 2013; Yano et al., 2016). In this study, we developed a Pairwise Comparison Analysis for Multiple Pool-seq, which is a powerful tool for cost-effective, rapid and efficient identification of genes related to anthocyanin biosynthesis in rice pericarp.

## Resuilts

### Phenotypic differences in second filial generation population and construction of bulks

In November 2016, when the rice grain was fully ripe and the moisture content reached 12.5%, grains were randomly selected and hulled, and then the seed coat color of F2 population was identified. The 381 lines developed from Huanghuazhan (HHZ) and Donglanmomi (DLMM) were divided into colored groups(including black, partial black and brown) and white group, and the number ratio was 285:96, which accorded with the separation rule of 3:1 (χ^2^ = 0.0079, P > 0.05). In November 2017, the number ratio of colored group to white group was 601:195 for 796 F2 lines, which accorded with the separation comparison of 3:1 (χ^2^ = 0.11, P > 0.05). Subsequently, the anthocyanin content of 601 colored pericarps and 30 white pericarps was measured by high performance liquid chromatography (HPLC). The results showed that the seed coats color was mainly caused by cyanidin content, and the content of cyanidin varied from 0.45 ug/g to 1616.03 ug/g. According to the seed coat color and anthocyanin content of 596 lines, the population was divided into four subgroups, and then 30 lines from each subgroup were selected to develop four bulks: white bulk (W), black bulk (B1), partial black bulk (B2) and brown bulk (B3)(Figure.1).

**Figure 1.**
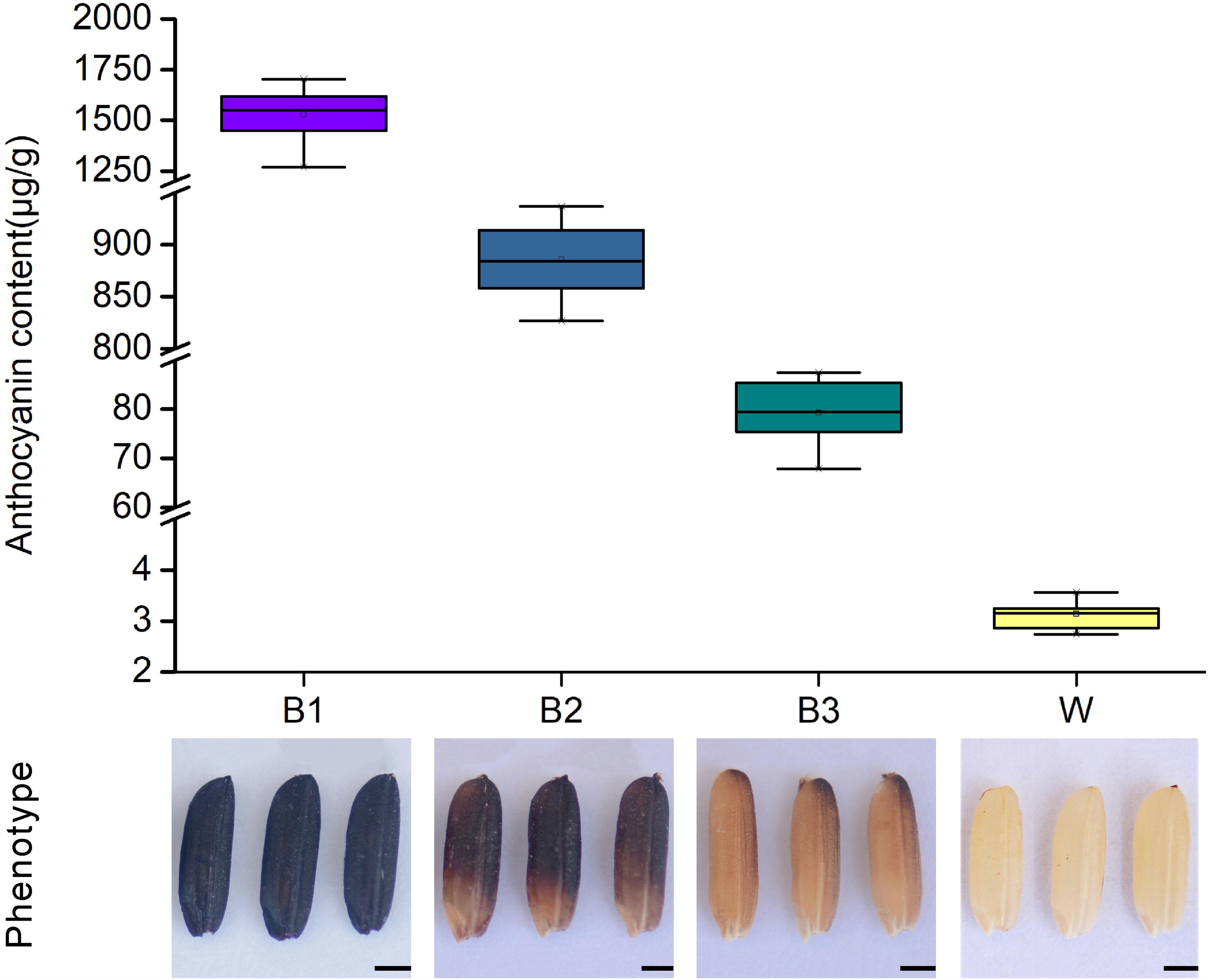
The phenotype and anthocyanin content in four pools of F2 population. Black pericarp pool (B1), partial peicarp black pool (B2), brown pericarp pool (B3), white pericarp pool (W). Scale bars, 1.2 mm.

### Mapping of reads to the reference genome and identifition of high quality SNPs

HHZ, DLMM, W, B1, B2 and B3 were sequenced using Illumina HiSeq X Ten sequencing platform. After filtration, the total base number of 6 bulks was 161.48 Gb, of which Huanghuazhan was 36.55 Gb, Donglanmomi was 39.63 Gb, W was 19.66 Gb, B1 was 22.11 Gb, B2 was 21.64 Gb and B3 was 21.89 Gb. The reads obtained by resequencing were aligned to the reference genome (http://rice.plantbiology.msu.edu/index.shtml) by BWA software. The alignment rates of Huanghuazhan, Donglanmomi, W, B1, B2 and B3 were 98.40%, 98.73%, 98.54%, 98.48%, 98.62% and 98.57 respectively. The average sequencing depth of 6 samples was 64.76x and the coverage of at least 1x was 94.98% (Table S1). According to the coverage depth of 12 chromosomes in rice (Figure. S1), the genomes of 6 samples were evenly covered and the sequencing randomness was good.

GATK(Mckenna et al., 2010A) was used to detect the SNPs between the bulks and the reference genome. A total of 2,427,862 SNPs were obtained from 6 bulks, including Huanghuazhan 2,123,666 SNPs, Donglanmomi 576,236 SNPs, W bulk 2,265,539 SNPs, B1 bulk 2,238,962 SNPs, B2 bulk 2,264,753 SNPs and B3 bulk 2,301,734 SNPs (Table S2). SnpEff (Cingolani et al., 2012A) was used to annotate and predict the variation effects of these SNPs. The results showed that most of the variation sites were located in upstream region and downstream region of the genes, which may affect the gene function.

Before association analysis, SNP was filtered. The criteria were as follows: firstly, SNPs with multiple genotypes were filtered out; secondly, SNPs with read depth less than 4 were filtered out; thirdly, SNPs with identical genotypes between bulks and SNPs of recessive bulk which were not from recessive parent were filtered out. Finally, W-B1 bulk obtained 1,668,781 high-quality SNPs, W-B2 bulk 1,674,742 high-quality SNPs, W-B3 bulk 1,683,759 high-quality SNPs, B1-B2 bulk 1,669,167 high-quality SNPs, B1-B3 1,680,364 high-quality SNPs, B2-B3 bulk 1,688,944 high-quality SNPs (Table S3).

### Anthocyanin biosynthesis related genomic region in rice pericarp

To identify the candidate genomic regions responsible for anthocyanin biosynthesis in rice pericarp, we compared the SNP-index(Takagi et al., 2013C) between different bulks. Distance method was used to fit the Δ(SNP-index). The distribution of SNP-index and Δ(SNP-index) (Figure. 2) of W–B1 showed that the regions showing significant association with anthocyanin biosynthesis in rice pericarp were mapped to 26.59–30.92 Mb interval on chromosome 2 (Table S4), 8.76–10.07 Mb interval on chromosome 3 (Table S5), and 22.76–32.12 Mb interval on chromosome 4 (Table S6).

**Figure 2.**
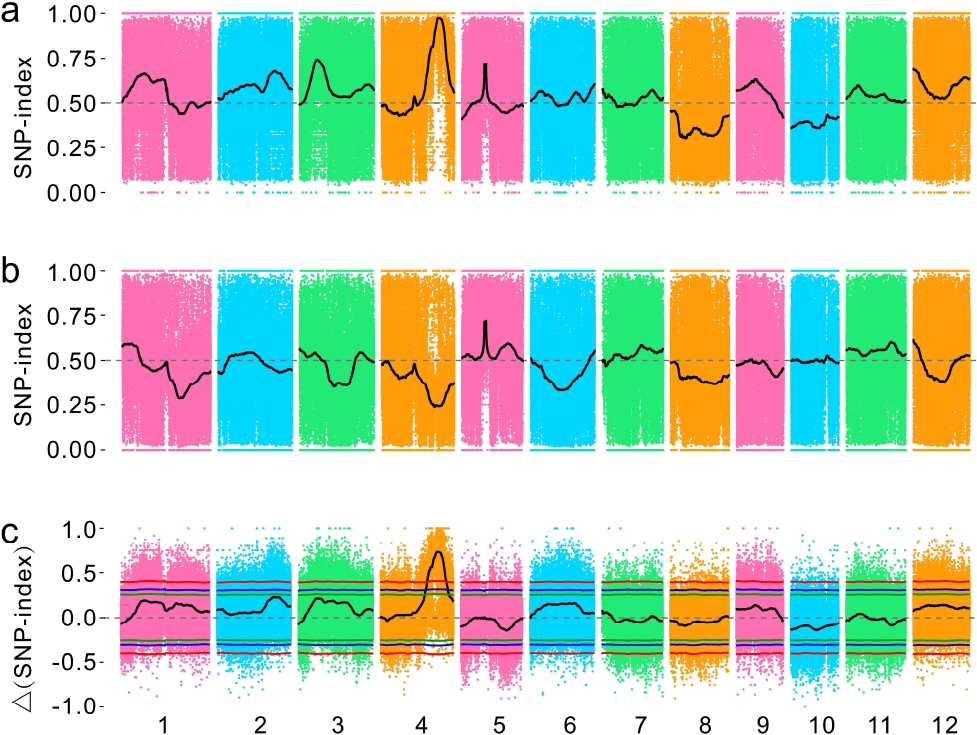
The PCAMP approach for mapping genomic regions controlling anthocyanin biosynthesisbetween W and B1. SNP-index plots of W (a) and B1 (b), Δ (SNP-index) plot (c) of chromosome. The y-axis is the name of the chromosome, colored dots represent the calculated SNP-index, and the black line is the fitted SNP-index. The green line, blue line and red line represent the threshold of 90%, 95% and 99% confidence interval, respectively.

The distribution of SNP-index and Δ(SNP-index) (Figure. 3) of W–B1 showed that the regions showing significant association with anthocyanin biosynthesis in rice pericarp were mapped in 32.34–34.95 Mb interval on chromosome 3 (Table S7), 25.17–32.21 Mb interval on chromosome 4 (Table S8), and 2.76–5.46 Mb interval on chromosome 12 (Table S9).

**Figure 3.**
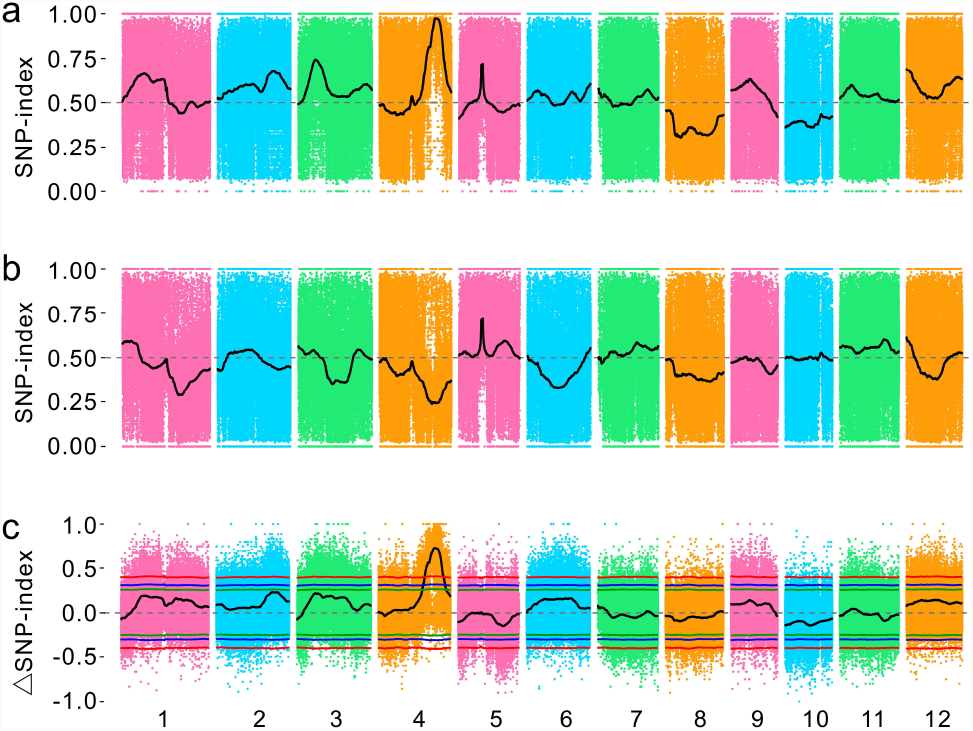
The PCAMP approach for mapping genomic regions controlling anthocyanin biosynthesisbetween W and B2. SNP-index plots of W (a) and B2 (b), Δ (SNP-index) plot (c) of chromosome. The y-axis is the name of the chromosome, colored dots represent the calculated SNP-index, and the black line is the fitted SNP-index. The green line, blue line and red line represent the threshold of 90%, 95% and 99% confidence interval, respectively.

The distribution SNP-index and Δ(SNP-index) (Figure. 4) of W–B3 showed that the regions showing significant association with the genes involved in anthocyanin biosynthesis in rice pericarp were mapped to 25.15–31.14 Mb interval on chromosome 4 (Table S10) and 7.11–9.80 Mb interval on chromosome 9 (Table S11).

**Figure 4.**
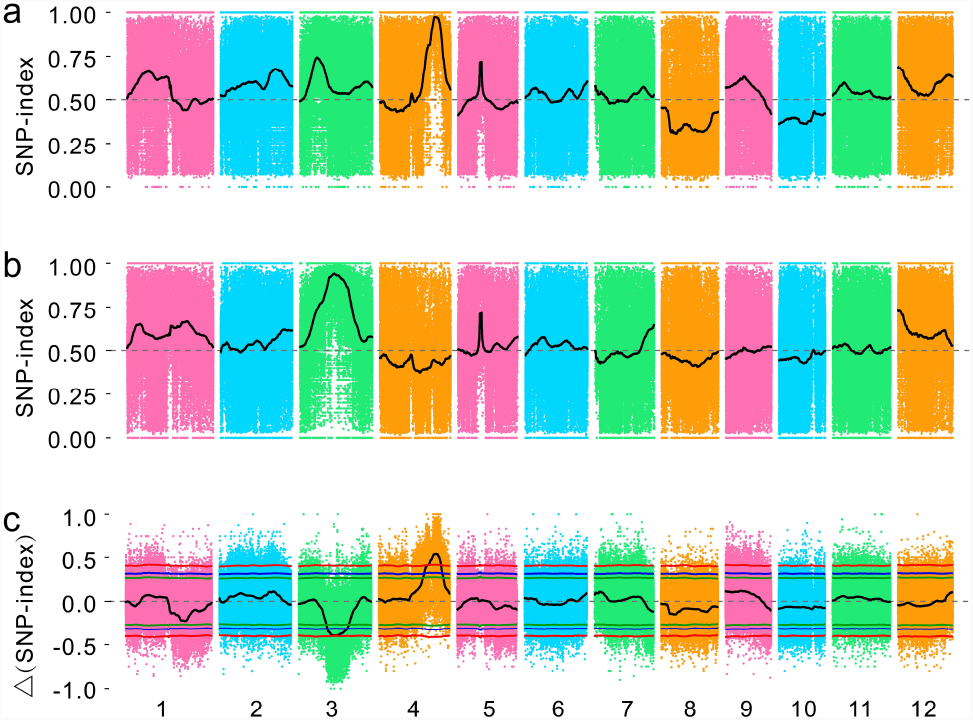
The PCAMP approach for mapping genomic regions controlling anthocyanin biosynthesisbetween W and B3. SNP-index plots of W (a) and B3 (b), Δ (SNP-index) plot (c) of chromosome. The y-axis is the name of the chromosome, colored dots represent the calculated SNP-index, and the black line is the fitted SNP-index. The green line, blue line and red line represent the threshold of 90%, 95% and 99% confidence interval, respectively.

The distribution of SNP-index and Δ(SNP-index) (Figure. 5) of B1–B2 showed that the regions showing significant association with genes involved in anthocyanin biosynthesis in rice pericarp were mapped to the 13.30–24.65 Mb interval on chromosome 3 (Table S12) and 8.09–10.92 Mb, 10.94–1.37 Mb, 11.97–12.13 Mb, 12.24–12.25 Mb, 12.28–12.34 Mb, 12.36–13.06 Mb, 14.07–16.96 Mb, 16.99–17.14 Mb intervals on chromosome 6 (Table S13).

**Figure 5.**
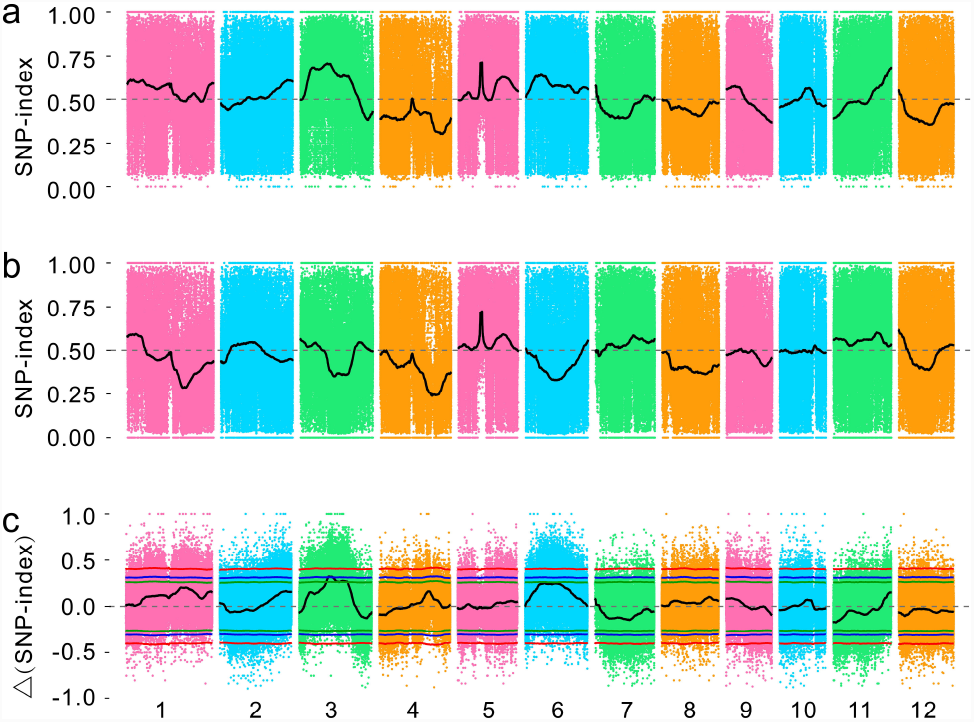
The PCAMP approach for mapping genomic regions controlling anthocyanin biosynthesisbetween B2 and B1. SNP-index plots of B2 (a) and B1 (b), Δ (SNP-index) plot (c) of chromosome. The y-axis is the name of the chromosome, colored dots represent the calculated SNP-index, and the black line is the fitted SNP-index. The green line, blue line and red line represent the threshold of 90%, 95% and 99% confidence interval, respectively.

The distribution of SNP-index and Δ(SNP-index) (Figure. 6) of B1–B3 showed that the regions showing significant association with genes involved in anthocyanin biosynthesis in rice pericarp were mapped to 26.57–31.55 Mb interval on chromosome 1 (Table S14) and 13.58–25.11 Mb interval on chromosome 3 (Table S15).

**Figure 6.**
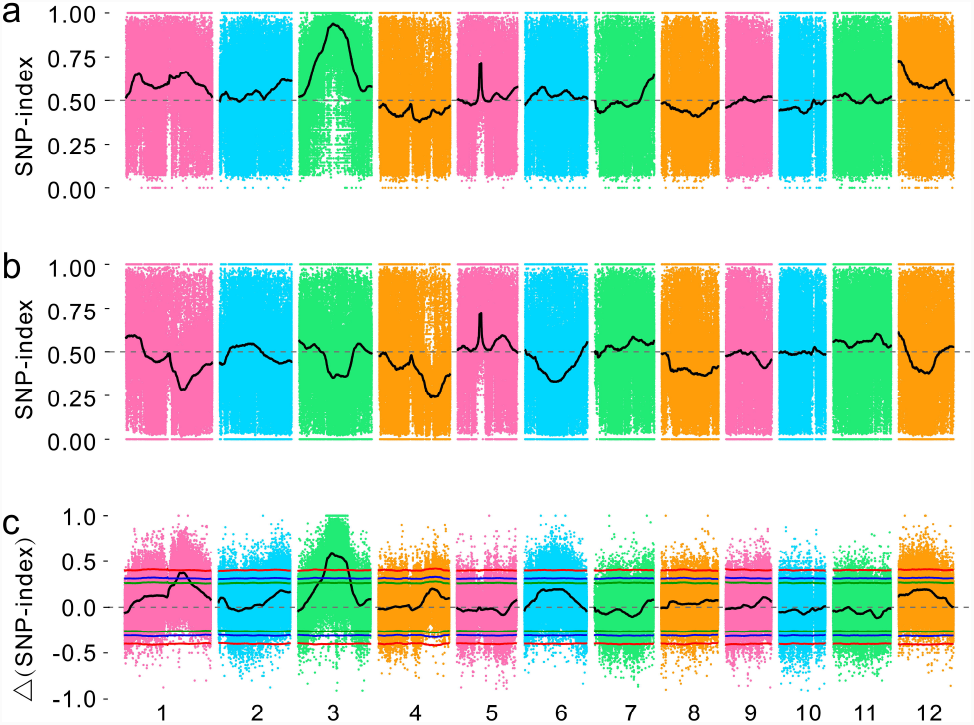
The PCAMP approach for mapping genomic regions controlling anthocyanin biosynthesisbetween B3 and B1. SNP-index plots of B3 (a) and B1 (b), Δ (SNP-index) plot (c) of chromosome. The y-axis is the name of the chromosome, colored dots represent the calculated SNP-index, and the black line is the fitted SNP-index. The green line, blue line and red line represent the threshold of 90%, 95% and 99% confidence interval, respectively.

The distribution of SNP-index and Δ(SNP-index) (Figure. 7) of B2–B3 showed that the regions showing significant association with genes involved in anthocyanin biosynthesis in rice pericarp were mapped in 17.22–20.86 Mb and 20.94–21.02 Mb intervals on chromosome 3 (Table S16) and 1.64–16.96 Mb interval on chromosome 12 (Table S17).

**Figure 7.**
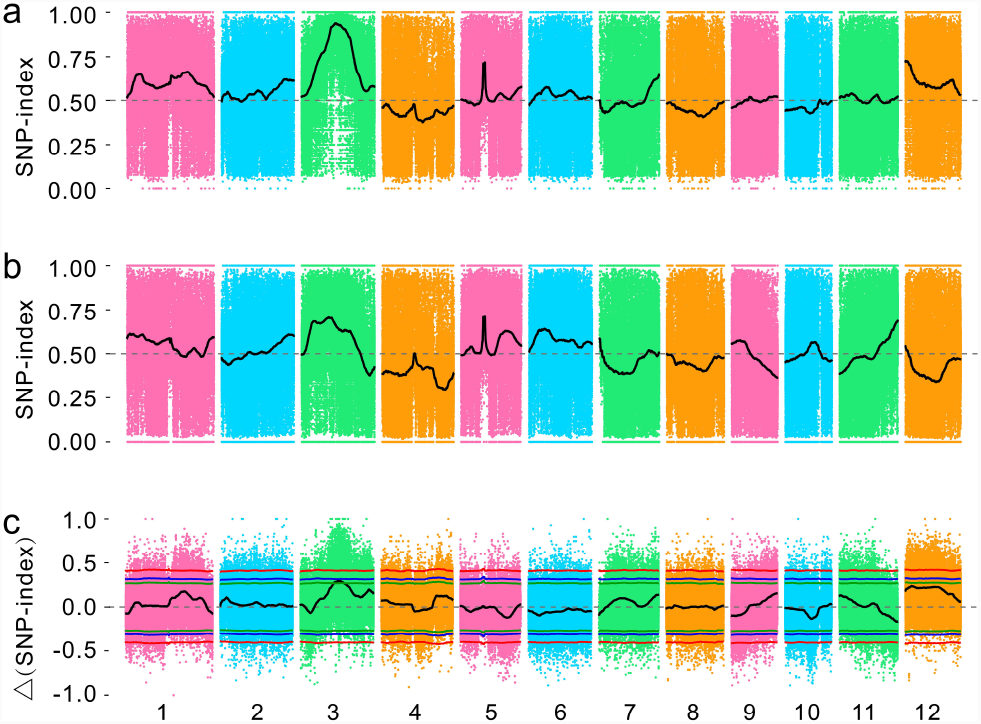
The PCAMP approach for mapping genomic regions controlling anthocyanin biosynthesisbetween B3 and B2. SNP-index plots of B3 (a) and B2 (b), Δ(SNP-index) plot (c) of chromosome. The y-axis is the name of the chromosome, colored dots represent the calculated SNP-index, and the black line is the fitted SNP-index. The green line, blue line and red line represent the threshold of 90%, 95% and 99% confidence interval, respectively.

For the candidate genomic regions with overlapping physical positions on the same chromosome, the intersection regions were selected as the final candidate regions. Therefore, the regions showing significant association with genes involved in anthocyanin biosynthesis in rice pericarp were shown in Table S18.

### Candidate genes for anthocyanin biosynthesis in rice pericap

We classified the candidate genomic regions into two groups: (I) the regions that was adjacent to or contained the cloned genes related to anthocyanin biosynthesis genes; and (II) the remaining candidate regions.

(I): The *Rd* is located at 1.19 Mb upstream of the candidate region on chromosome 1. Furukawa *et al.*(Furukawa et al., 2007) found that *Rd* was involved in the proanthocyanidin biosynthesis of rice pericarp. The expression level of *Rd* between HHZ and DLMM was significantly different (Figure. 8a). The sequences of HHZ and DLMM were amplified with PCR primer AB003495(Konishi et al., 2008) and the products were sequenced (Table S19). The 43rd base of the second exon of the *Rd* of HHZ was changed from C to A. This change belongs to the *Rd2* mutant genotype(Konishi et al., 2008) (Figure. 8b).

**Figure 8.**
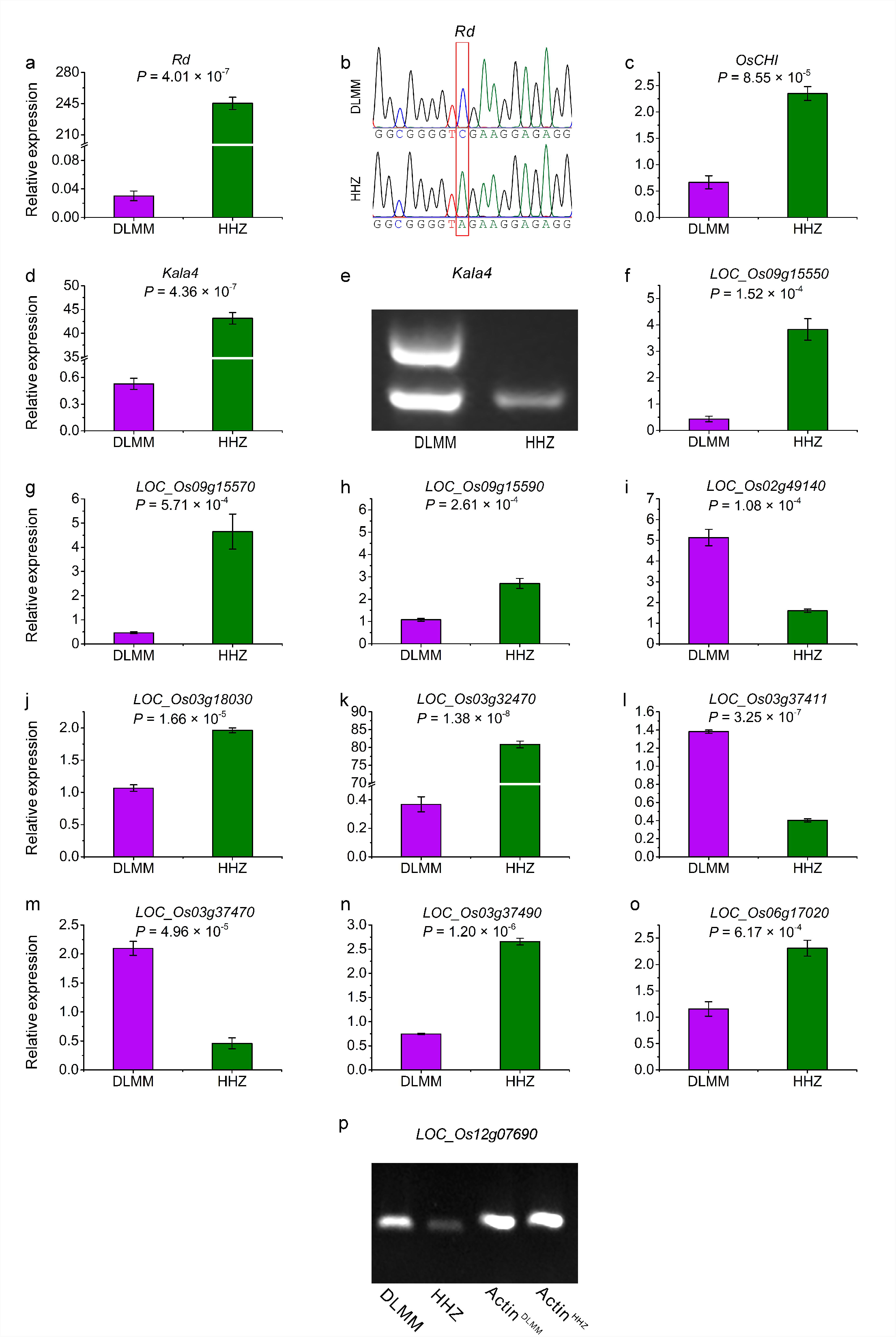
The candidate genes of anthocyanin biosynthesis in rice pericarp. The expression level of 13 genes through qPCR (a, c, d, f–q). DNA sequencing of Rd and the 43rd base of the second exon is mutated from C to A (b). Amplification of Kala4 using functional primers (e). The expression level of *LOC_Os12g07690* through RT-PCR (p).

Druka *et al*.(Druka et al., 2003) characterized the anthocyanin synthesis gene *OsCHI* based on the maize homolog gene *Cfi* within the 32.34–34.95 Mb interval on chromosome 3. Using golden hull and internode mutants of rice, Hong *et al*.(Hong et al., 2012) cloned *OsCHI*, a key gene involved in flavonoid metabolic pathway. The expression levels of *OsCHI* between HHZ and DLMM were significantly different (Figure. 8c).

Hu *et al.*(Hu et al., 1996) identified the *Ra* gene using the maize homologous sequences within 25.17–31.14 Mb interval on chromosome 4, *Ra* gene encodes the basic helix-loop-helix (bHLH) transcription factor, which plays a regulatory role in the anthocyanin synthesis pathway. Subsequently, Hu *et al.*(Hu et al., 2000) indicated that *Ra* was consisted of *Ra1* and *Ra2*. Recently, Oikawa *et al*.(Oikawa et al., 2015) successfully cloned *Kala4*, a key gene responsible for anthocyanin accumulation in rice pericarp, which was found to be the same gene as *Ra2*. The expression levels of *Kala4* between HHZ and DLMM were significantly different (Figure. 8d). The DNA samples of HHZ and DLMM were amplified by functional primers (Table S19). The results showed that the promoter region of *Kala4* in DLMM had a genomic fragment inserted (Figure. 8e), and this change was the causes of generation of the black rice traits(Oikawa et al., 2015).

At the 0.24 Mb upstream of the candidate region on chromosome 9, Shao *et al.*(Shao et al., 2012) found that *inhibitor for brown furrows1* 1 (*IBF1*) might be involved in the accumulation of anthocyanins in hull, but there was no significant difference in *IBF1* expression level between HHZ and DLMM. However, the expression levels of *LOC_Os09g15550, LOC_Os09g15570* and *LOC_Os09g15590* (Figure. 8f–h) between HHZ and DLMM were significantly different. These three genes are similar to the *IBF1* gene and they all contain F-box domain.

(II): There are 61 SNPs with Δ(SNP-index)>=0.67 in 26.59–30.92 Mb intervals on chromosome 2. They included a homozygous variant site of Δ(SNP-index)=1 (Table S20). The expression levels of *LOC_Os02g49140* between HHZ and DLMM were significantly different (Figure. 8i), and this gene encodes glycosyltransferase. Previous studies have shown that glycosylation modification of anthocyanin affects its stability in cells(Fukuchi-mizutani et al., 2003).

Within the 8.76–10.07 Mb region on chromosome 3, there are 24 SNPs with Δ(SNP-index)>=0.67, and 2 homozygous variant loci with Δ(SNP-index)=1 (Table S21). The expression levels of *LOC_Os03g18030* between HHZ and DLMM were significantly different (Figure. 8j). This gene encodes leucoanthocyanidin dioxygenase, a key enzyme involved in anthocyanin biosynthesis pathway in plants(Saito et al., 1999).

The region of 17.22–21.02 Mb interval on chromosome 3 is consistent with that reported in previous studies(Yang et al., 2018). There were 4620 SNPs with Δ(SNP-index)>=0.67, including 69 homozygous variant sites with Δ(SNP-index)=1 (Table S22). The expression levels of *LOC_Os03g32470, LOC_Os03g37411, LOC_Os03g37470* and *LOC_Os03g37490* (Figure. 8k–n) between HHZ and DLMM were significantly different. *LOC_Os03g32470* encodes leucoanthocyanidin dioxygenase, which catalyzes the oxidative dehydration of leucocyanidins to form the anthocyanins(Saito et al., 1999). The other three genes encode MATE efflux family protein. *LOC_Os03g37411* and *LOC_Os03g37490* are highly homologous to *AtTT12* of *Arabidopsis thaliana* (Figure. S2). In *Arabidopsis, AtTT12* is involved in the transport of anthocyanins or proanthocyanidins to vacuoles(Marinova et al., 2007; Debeaujon et al., 2001). There were 96 SNPs with Δ(SNP-index)>=0.67 in 8.09–17.14 Mb interval on chromosome 6, including two homozygous mutation sites of Δ(SNP-index)=1 (Table S23). The expression levels of *LOC_Os06g17020* between HHZ and DLMM were significantly different (Figure. 8o). *LOC_Os06g17020* encodes anthocyanin 3-O-beta-glucosyltransferase, a key enzyme catalyzing the oxidation of unstable anthocyanidins into anthocyanins(Fukuchi-mizutani et al., 2003).

There were 40 SNPs with Δ(SNP-index)>=0.67 in 2.76–5.46 Mb interval on chromosome 12, including a homozygous variation site of Δ(SNP-index)=1 (Table S24). The expression levels of *LOC_Os12g07690* between HHZ and DLMM were significantly different (Figure. 8p). The function of *LOC_Os12g07690* which encodes chalcone synthase is related to flavonoid biosynthesis (https://phytozome.jgi.doe.gov/pz/portal.html#).

## Discussion

The SHOREmap (Schneeberger et al., 2009), NGM (Austin et al., 2011), Mutmap)(Abe et al., 2012), and QTL-seq (Takagi et al., 2013C) were developed based on the combination of high throughput sequencing and BSA technology. However, SHOREmap technology needs a large number of samples, which seriously limits the application; the mapping population used by NGM is usually recessive homozygous population, which is applicable to recessive homozygous mutation phenotype; although the bulk sequencing for QTL analysis pipeline developed by Magwene et al. [53] can detect multiple genes, even minor effect genes, affecting the target traits, but the algorithm is complex and not practical. MutMap is an effective method for studying important traits of plants and animals with its advantages of high efficiency and accuracy. However, MutMap is mainly applicable to quality traits controlled by single gene, and not suitable for some quantitative traits controlled by multiple genes. QTL-seq and its derivatives technologies have the advantages of low cost, short period and accurate mapping. It has been widely used in the mapping of quality and quantitative traits in plants. However, only 1 or 2 major effect genes related to target traits can be identified. It is often helpless for complex traits controlled by multiple genes.

GWAS can be used to map complex plant traits and quickly identify genetic loci controlling target traits. However, GWAS is applicable to natural population with a large sample size leading to high cost and it is also difficult to detect the rare mutations and minor effective genes (Platt et al., 2010; Farlow, 2013; Yano et al., 2016). Family-based QTL mapping plays an important role in the mapping of complex plant traits. This method can identify major and minor loci related to target traits, but the population size is large and multiple generations required to develop pedigrees(Mitchell-olds, 2010).

PCAMP technology divides the segregated population into n subgroups (n ≥ 3) according to the phenotype of the target traits. For every subgroup, the DNA of individual selected from the subgroups was pooled. High throughput sequencing technology is used to sequence the bulk. Then, combination of every two bulks is respectively compared to obtain target traits related QTLs. Using PCAMP technology, we successfully identified 10 candidate intervals related to anthocyanin synthesis in rice seed coat. Among the 10 candidate intervals, *Rd*(Furukawa et al., 2007), *OsCHI* (Hong et al., 2012; Druka et al., 2003), and *Kala4*(Oikawa et al., 2015) have been cloned, and two regions overlapped with that identified by provious GWAS study (Yang et al., 2018). These results validated the reliability of the method.

The candidate regions on chromosome 4 related to anthocyanin synthesis in rice seed coat could be detected by PCAMP in W-B1, W-B2 and W-B3 combinations. Finally, we selected three candidate regions that are shared for the real candidate regions, and the candidate gene *Kala4* was just located in this region(Oikawa et al., 2015). Likewise, the overlapping regions of candidate regions B1–B2, B2–B3, and B1–B3 on chromosome 3 were used and the candidate region was finally determined to be 17.22–20.86 Mb interval and 20.94–21.02 Mb, which are consistent with our previous study(Yang et al., 2018). The final interval 2.76–5.46 Mb interval on chromosome 12 is only 17.7% of the maximum candidate interval. Therefore, the candidate region of the target gene can be narrowed down by using the PCAMP.

Kala4 can be identified by PCAMP in the combinations of three colored pools and white pool, which plays an important role in anthocyanin synthesis in rice seed coat. However, when the gene kala4 exists, anthocyanin can not be synthesized regardless of the genotype of other genes, which is consistent with the previous report (Maeda et al., 2014). In the combinations of three colored pools, 17.22-20.86 Mb and 20.94-21.02 Mb of chromosome 3 were identified, which indicated that there were at least two key genes related to anthocyanin synthesis. In the previous study, we used 419 Core Germplasms of rice landraces in Guangxi for GWAS and identified that this region was an important candidate region for anthocyanin synthesis genes in rice pericarpdy (Yang et al., 2018). The *Rd* was located at 1.19 Mb upstream of the candidate region. What is the reason for this? In this study, we found that the four pools were composed of DNA samples of 30 individuals, and their mean sequencing depth was 64.76×. Therefore, this was not due to the number of individuals in pools and the depth of sequencing(James et al., 2013). Subsequently, we analyzed the distribution of SNPs on 12 chromosomes between HHZ and DLMM (Figure. S3). The results showed that the number of SNPs in the genomic region nearby *Rd* was greatly reduced (Figure. 9). Huang *et al.*(Huang et al., 2012) found that the nucleotide polymorphism of selective sweep regions (22.4–23.3Mb) on chromosome 1 was low. *Rd* is the domestication-related gene(Konishi et al., 2008), thus, the false positive result may be resulted from a decrease in nucleotide polymorphism within this genomic region.

**Figure 9.**
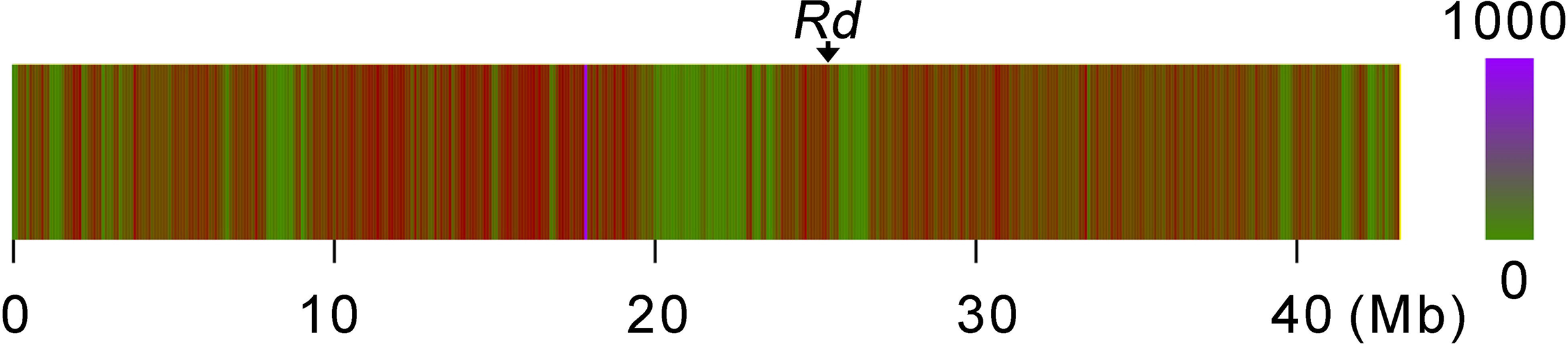
The *Rd* is located within the selective sweep regions on chromosome 1.

Anthocyanins are important pigments for seed plants. They are catalyzed by a series of enzymes encoded by structural genes and transported to vacuoles or other parts for storage. In the anthocyanin synthesis pathway, the modifications on downstream pathways are highly species-specific. After the unstable leucoanthocyanidins synthesized into stable anthocyanidins, they undergo glycosidation to become stable anthocyanins, and this step is catalyzed by glycosyltransferase (Fukuchi-mizutani et al., 2003). *LOC_Os02g49140* encodes Glycosyltransferase, and *LOC_Os06g17020* encodes anthocyanin 3-O-beta-glucosyltransferase.

Anthocyanidin synthase (ANS) is one of the four dioxygenases (DOX) involved in flavonoid biosynthetic pathway. It catalyzes the oxidation of colorless anthocyanins to produce colored anthocyanins. Reddy et al. (Reddy et al., 2007) used homologous cloning to find the location of rice *ANS* gene on the short arm end of the chromosome 1. Overexpression of this gene leads to accumulation of a mixture of flavonoids and anthocyanins. *ANS* is mainly expressed in rice leaves and seeds (Kim et al., 2007). Both *LOC_Os03g18030* and *LOC_Os03g32470* encode for leucoanthocyanidin dioxygenase, but their expression levels are significantly differed between HHZ seeds and DLMM seeds, especially *LOC_Os03g32470*.

After the synthesis of anthocyanin, if it stays in cytoplasm, its high biochemical reactivity will cause toxicity to cells, and anthocyanin itself may also be oxidatively denatured during the reaction (Zhao j, 2010). There are mainly three types of vacuolar transport for anthocyanins, one of which is to transport anthocyanins into vacuole via the MATE transporter on the tonoplast. This process requires the H^+^ concentration gradient produced by H^+^-ATPase proton pump (Appelhagen et al., 2015). *LOC_Os03g37411, LOC_Os03g37470*, and *LOC_Os03g37490* all encode for MATE efflux family proteins. *LOC_Os03g37411* and *LOC_Os03g37490* are highly homologous to *AtTT12* gene in *Arabidopsis thaliana*. In addition, *TT12* also plays an important role in the flavonoid metabolism of plants, such as rape (Chai et al., 2009) and cotton (Gao et al., 2016).

F-box protein amily is widely distributed proteins in eukaryotes. They contain F-box domains and participate in the regulation of many growth and development processes in plants. Studies have shown that there are 687 F-box proteins in rice (Jain et al., 2007). *BF1* encodes for a F-box protein rich in Kelch repeats. This gene is a negative regulator in the flavonoid regulatory pathway, and it inhibits the accumulation of total flavonoids and anthocyanins in rice hulls (Shao et al., 2012). *LOC_Os09g15550, LOC_Os09g15570* and *LOC_Os09g15590* all encode for F-box domain containing proteins.

Chalcone synthase is the first enzyme in the biosynthesis pathway of flavonoid secondary metabolites. It catalyzes the formation of naringenin chalcone by coumaroyl-CoA and malonyl-CoA, providing starting materials for the biosynthesis of flavonoid secondary metabolites. OsCHS1 and OsCHS2 have been identified in rice by homologous cloning. The results of yeast two-hybrid analysis indicate that OsCHS1 can interact with *OsF3H, OsF3’H, OsDFR* and *OsANS1* (Shih et al., 2008). High-throughput sequencing was used to analyze the induced mutant Red-1, and the result showed that the *BGIOSGA033874* gene, which was related to flavonoid synthesis pathway in rice, had the same locus as *OsCHS1* (Cheng et al., 2014). *LOC_Os12g07690* encodes for Chalcone synthase, which may be involved in flavonoid biosynthesis.

## Materials and methods

### Plant materials

DLMM (rank No: C133) is the core germplasm of rice landraces in Guangxi(Yang et al., 2018). Its pericarp is black and the anthocyanin content is high. The anthocyanin is mainly cyanidin, accounting for 98.4% of total anthocyanin content. HHZ pericarp is white and almost without anthocyanins. The target traits of the parent materials vary greatly (Figure. S4). The F_1_ and F_2_ populations were constructed by crossing and backcrossing with HHZ and DLMM, respectively. The two parents, F_1_ and F_2_ populations were planted in Nanning Experimental Field (22.85 ºN, 108.26 ºE, Guangxi, China) from July to November, 2017.

### Determination of pericarp color and anthocyanin content

After the rice seeds were fully matured and dried to a water content of 12.5%, the whole kernels were randomly selected for shelling, and the pericarp colors of the individuals in the two parents and the F_2_ population were identified according to the method described by Han & Wei(Han et al., 2006). The content of anthocyanin in brown rice was determined by HPLC method(Beta et al., 2009), which mainly included geranium pigment, morning glory pigment, delphinidin pigment, peony pigment, cyanidin and mallow pigment.

### Construction of Genomic DNA pools

In the late 2017 season, 1–2 young leaves at the seedling stage of each individual plant in the F_2_ population were harvested and then stored in a −20°C refrigerator for later use. According to the pericarp color and anthocyanin content of each individual grain, the isolated population was divided into four subpopulations, namely black pericarp subpopulation, partial peicarp black subpopulation, brown pericarp subpopulation, and white pericarp subpopulation. Then, 30 individual plants were selected from each subpopulation and DNA was extracted by cetyltrimethylammonium bromide (CTAB) method(Murray et al., 1980) to form a DNA pool.

### Illumina sequencing

After concentrations of DNA samples of 2 parents and 4 mixed pools were measured, the qualified DNA samples were randomly broken into 350 bp fragments by ultrasonic disruption, and the DNA fragments were terminated by end repair at the 3’ end with ploy A, plus sequencing linker, purification, and PCR amplification. A sequencing library was constructed. After being passed the quality check, the library was sequenced by Illumina HiSeq X Ten.

### Alignment with reference genome and SNP annotation

The reads obtained by re-sequencing need to be relocated to the rice reference genome (http://rice.plantbiology.msu.edu/) for subsequent variation analysis. The short sequences obtained by sequencing were aligned with the reference genome using BWA software(Li et al., 2009). After reading the reference genome, reads can count the coverage of the base on the reference genome. SNPs were detected using GATK software(Mckenna et al., 2010B). According to the positioning results of Clean Reads in the reference genome, Picard was used for De-duplication (Mark Duplicates), GATK was used for Local Realignment, Base Recalibration, etc., and then GATK was used for single core. After detection of single nucleotide polymorphism (SNP) and filtering, a final set of SNP sites was obtained. Annotation variation and predictive variation effects were performed using SnpEff software(Cingolani et al., 2012B). Based on the position of the variant site on the reference genome and the position information of the gene on the reference genome, the region harboring the mutation site in the genome and the effect of the mutation were obtained.

### SNP-index analysis

First, the SNPs are filtered as follows: (1) SNP loci were filtered out with multiple genotypes; (2) SNP loci were filtered out with read support less than 4; and (3) SNPs were filtered out with consistent genotypes between pools. The loci and recessive pool genes are not SNP loci from recessive parents. Using the SNP data of the two parents, the SNP-index(Takagi et al., 2013C) of each mixed pool were calculated, and the SNP-indexes of each mixed pool were compared (Figure. S5). The calculation formula is as follows:

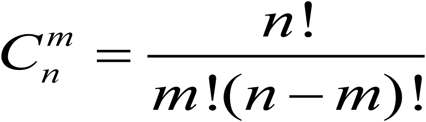

where n represents the number of all Pool-seqs, m (=2) represents any two mixed pools, and the candidate regions associated with the target traits are obtained by the SNP-index method.

### Identification of candidate gene

We selected a genomic region above the threshold corresponding to the 95% or 99% confidence interval as a candidate region for the target gene. The same candidate regions appear in the results of different ΔSNP-indexes, and the intersection of these regions was selected as the final candidate region. In the genomic candidate region, it is preferred to use the rice genome annotation site MSU-RGAP (http://rice.plantbiology.msu.edu/) to annotate the genes in the candidate region, and the genes related to plant anthocyanin synthesis(Zhu et al., 2015); secondly, the candidate region containing candidate gene or adjacent to the known gene related to anthocyanin synthesis in rice were taken as a candidate gene; finally, the homozygous variation site with a sequencing depth of not less than 4× and ΔSNP-index=1 was selected as a candidate gene. At the same time, by referring to the method of Yano *et al.*(Yano et al., 2016), all the polymorphisms in the candidate region were classified into three types. TypeIincluded the SNPs that were predicted to induce amino acid exchange or to change splicing junctions; TypeII included SNPs that were located at the 5’-flanking sequences of genes or the up-streams region containing promoter region; and Type III included SNPs that were located in a coding region but were not predicted to change an amino acid, an intron or a 3’-noncoding sequence.

### PCR reaction

25 μl 2×Es Taq Master Mix (Beijing, China), upstream primer: 1 μl of upstream primer, 1 μl of downstream primer, 1 μl of DNA template, 22 μl of dd H_2_O, using Gene AmpPCR system 9700 thermocycler (Perkin Elmer Cetus, USA). The reaction procedures were programmed as follows: 35 cycles of 94°C for 2 min, cycle (94°C for 30 s, 56°C for 30 s, and 72°C 2 kb/min), and finally extension at 72°C for 5 min.

### DNA sequencing

After the PCR product was recovered and homozygous, the ABI3730XL (Applied Biosystems, USA) sequencer was used for bidirectional sequencing with both positive and negative primers. DNA sequences were obtained and analyzed using ClustalX software.

### Agarose assay

The PCR product was assayed for the target fragment using a 1.5% agarose gel and visualized using a gel imaging analysis system (Biosens SC 850, Shanghai, China).

### Quantitative real time PCR

According to the dynamic variation of anthocyanin accumulation in the DLMM pericarp of rice (Figure. S6), the shelled kernels at 15 days after the flowering of DLMM and HHZ were selected, and then total RNA was extracted with Trizol method (Invitrogen, Carlsbad, CA, USA). Reverse transcription was performed by One-Step gDNA removal and cDNA synthesis superMix (TransGen, Beijing, China). The *Actin 1* (http://icg.big.ac.cn/index.php/Oryza_sativa) and *Os03g0234200*(Sun et al., 2018) were used as the internal reference genes for normalization of expression levels of candidate genes. The qRT-PCR primers are presented in Table S19. The qPCR reaction system was set as follows: 10 μl of SYBR Color qPCR Master Mix (2×), 0.5 l of forward primer (10 μM), 0.5 l of reverse primer (10 μM), 1 μl of template cDNA, 8 μl of dd H_2_O. Reaction procedures were programmed as follows: (1) 95°C, 5 min; (2) 95°C 15 s, 57°C 15s, 72°C 20s; (3) From 95 °C to 95°C, stop at 0.3°C for 1 s, read once. Data analysis was performed in accordance with Livak *et al*. (Livak et al., 2001).

## Data availability

The whole genome resequencing data were deposited in the Sequence Read Archive (SRA) database under the accession number SRR7903633, SRR7903634, SRR7903635, SRR7903636, SRR7903637, SRR7903638.

## Acknowledgements

Genome sequencing was conducted by the Biomarker Technologies, Beijing, China. This work was supported by the National Natural Science Foundation of China (31860371), National key R & D projects of China (2016YFD0100101-03), State Key Laboratory of Rice Biology (170102), Guangxi Natural Science Foundation of China (2015GXNSFAA139054, 2018GXNSFAA138124), Guangxi’s Ministry of Science and Technology (AB16380117).

## Author contributions

X.H.Y., D.T.L. and G.F.D. contributed to study design, X.Z.X. and F.Q.X. contributed to data analysis, Z.Q.Z., Y.Y.W. and Y.X.Z. contributed to determination of pericarp color and anthocyanin content, X.Z.X., H.F.L and G.X.D contributed to PCR, Gel electrophoresis and qRT-PCR, Y.Z. contributed to DNA sequencing, X.H.Y. and Z.Q.Z. wrote this manuscript, D.T.L. and G.F.D. revised the manuscript. All authors read and approve the paper.

## Supplementary Figures

**Figure S1** Genome-wide distribution of coverage depth. (a) Huanghuazhan, (b) Donglanmomi, (c) W, (d) B1, (e) B2, (f) B3. The x-axis is the chromosomal location, and y-axis is the value obtained by taking the logarithm (log2) of the depth of the corresponding position on the chromosome.

**Figure S2** Comparison of amino acid sequences in LOC_Os03g37411, LOC_Os03g3749 and homologs. Amino acid sequences of LOC_Os03g37411, LOC_Os03g3749 and homologs from Arabidopsis thaliana (AT3G21690, AT3G59030), Brachypodium stacei (Brast02G219700), Panicum hallii (Pahal.I02702), Populus trichocarpa (POPTR_0004s01620), Sorghum (Sb01g016110, Sb01g016140), Zea mays (GRMZM5G842695). (a) Multiple sequence alignment using ClustalX. (b) Phylogenetic tree constructed with neighbor joining analysis.

**Figure S3** Distribution of SNPs on 12 chromosomes of Huanghuazhan and Donglanmomi.

**Figure S4** The difference in the anthocyanin synthesis between a and b is on the pericarp. (a) Gross morphology of Huanghuazhan and Donglanmomi at the seedling stage, (b) The panicle of Huanghuazhan and Donglanmomi. Scale bars, 2 cm.

**Figure S5** Schematic diagram of PCAMP for identification of anthocyanin biosynthesis genes in rice. B1: black pericarp pool, B2: partial peicarp black pool, B3: brown pericarp pool, W: white pericarp pool.

**Figure S6** Anthocyanin accumulation in the DLMM grain. After flowering 2d, 3d, 5d, 9d, 11d, 13d, 15d, 17d, 19d, 21d, 23d, 25d. Scale bars, 3 mm.

## Supplementary Tables

**Table S1** Summary information for the genome resequencing statistics of six samples.

**Table S2** Summary information for the SNPs annotation of six samples.

**Table S3** The SNP statistics between any two pools.

**Table S4** Rice anthocyanin biosynthesis candidate genomic region on chromosome 2 between W and B1 at the 95% confidence interval.

**Table S5** Rice anthocyanin biosynthesis candidate genomic region on chromosome 3 between W and B1 at the 95% confidence interval.

**Table S6** Rice anthocyanin biosynthesis candidate genomic region on chromosome 4 between W and B1 at the 99% confidence interval.

**Table S7** Rice anthocyanin biosynthesis candidate genomic region on chromosome 3 betweenW and B2 at the 95% confidence interval.

**Table S8** Rice anthocyanin biosynthesis candidate genomic region on chromosome 4between W and B2 at the 99% confidence interval.

**Table S9** Rice anthocyanin biosynthesis candidate genomic region on chromosome 12 between W and B2 at the 95% confidence interval.

**Table S10** Rice anthocyanin biosynthesis candidate genomic region on chromosome 4 betweenW and B3 at the 99% confidence interval.

**Table S11** Rice anthocyanin biosynthesis candidate genomic region on chromosome 9 between W and B3 at the 95% confidence interval.

**Table S12** Rice anthocyanin biosynthesis candidate genomic region on chromosome 3 between B2 and B1 at the 95% confidence interval.

**Table S13** Rice anthocyanin biosynthesis candidate genomic region on chromosome 6 between B2 and B1 at the 95% confidence interval.

**Table S14** Rice anthocyanin biosynthesis candidate genomic region on chromosome 1 between B3 and B1 at the 95% confidence interval.

**Table S15** Rice anthocyanin biosynthesis candidate genomic region on chromosome 3 between B3 and B1 at the 99% confidence interval.

**Table S16** Rice anthocyanin biosynthesis candidate genomic region on chromosome 3 between B3 and B2 at the 99% confidence interval.

**Table S17** Rice anthocyanin biosynthesis candidate genomic region on chromosome 12 between B3 and B2 at the 95% confidence interval.

**Table S18** The final candidate genomic regions of anthocyanin biosynthesis in rice pericarp.

**Table S19** Primers used for this study.

**Table S20** List of 61 SNPs with ΔSNP-index≥0.67 in 26.59–30.92 Mb on chromosome 2.

**Table S21** List of 24 SNPs with ΔSNP-index≥0.67 in 8.76–10.07 Mb on chromosome 3.

**Table S22** List of 4620 SNPs with ΔSNP-index≥0.67 in 17.22–20.86 Mb and 20.94-21.02 Mb on chromosome 3.

**Table S23** List of 96 SNPs with ΔSNP-index≥0.67 in 8.09–17.14 Mb on chromosome 6.

**Table S24** List of 40 SNPs with ΔSNP-index≥0.67 in 2.76-5.46 Mb on chromosome 12.

