## Supplementary figures and images for "Pair-wise Comparison Analysis for Multiple Pool-seq: an efficient method identified anthocyanin biosynthesis genes in rice pericarp"

### Fig. S1.jpg

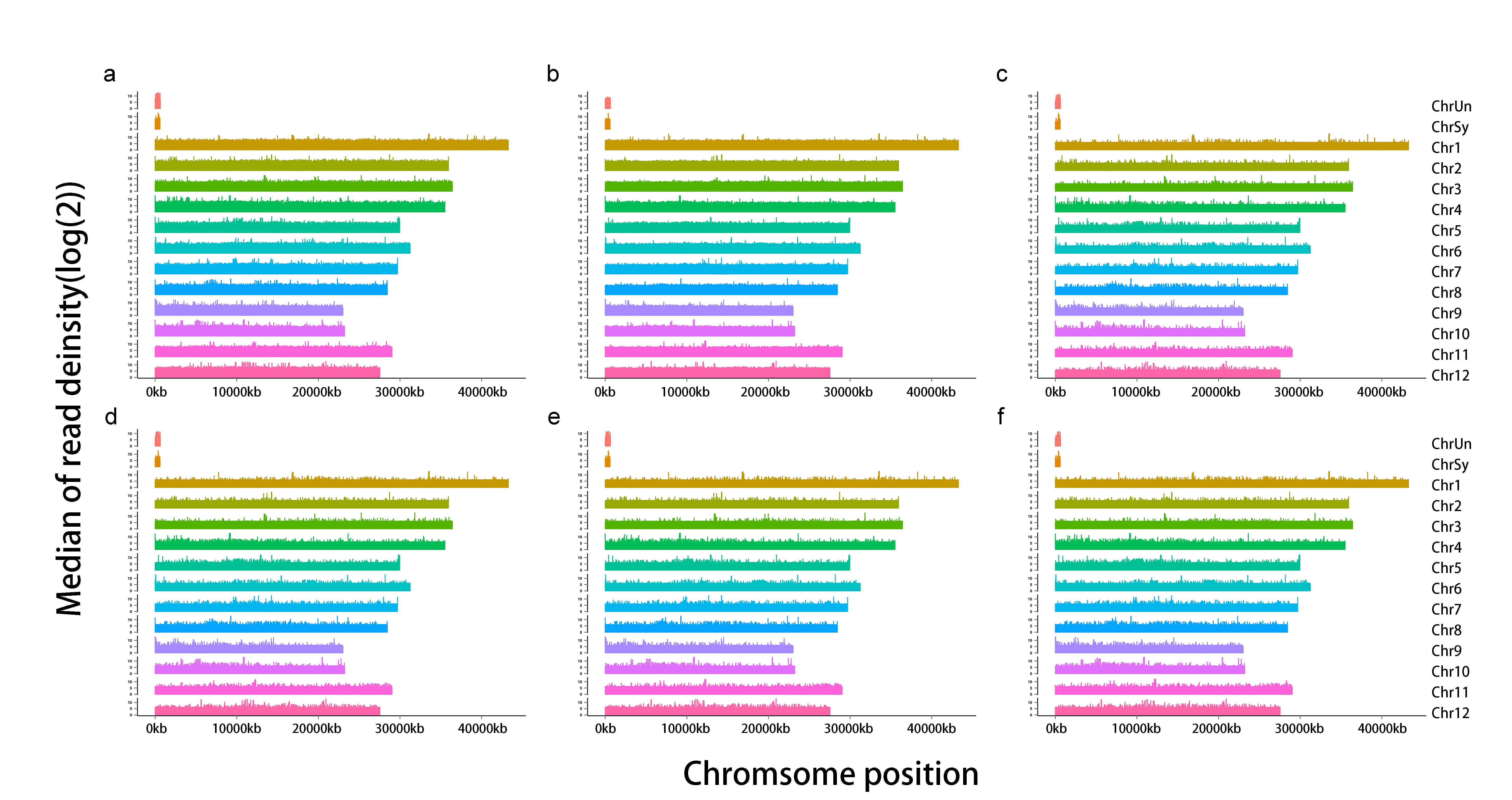

### Fig. S2.jpg

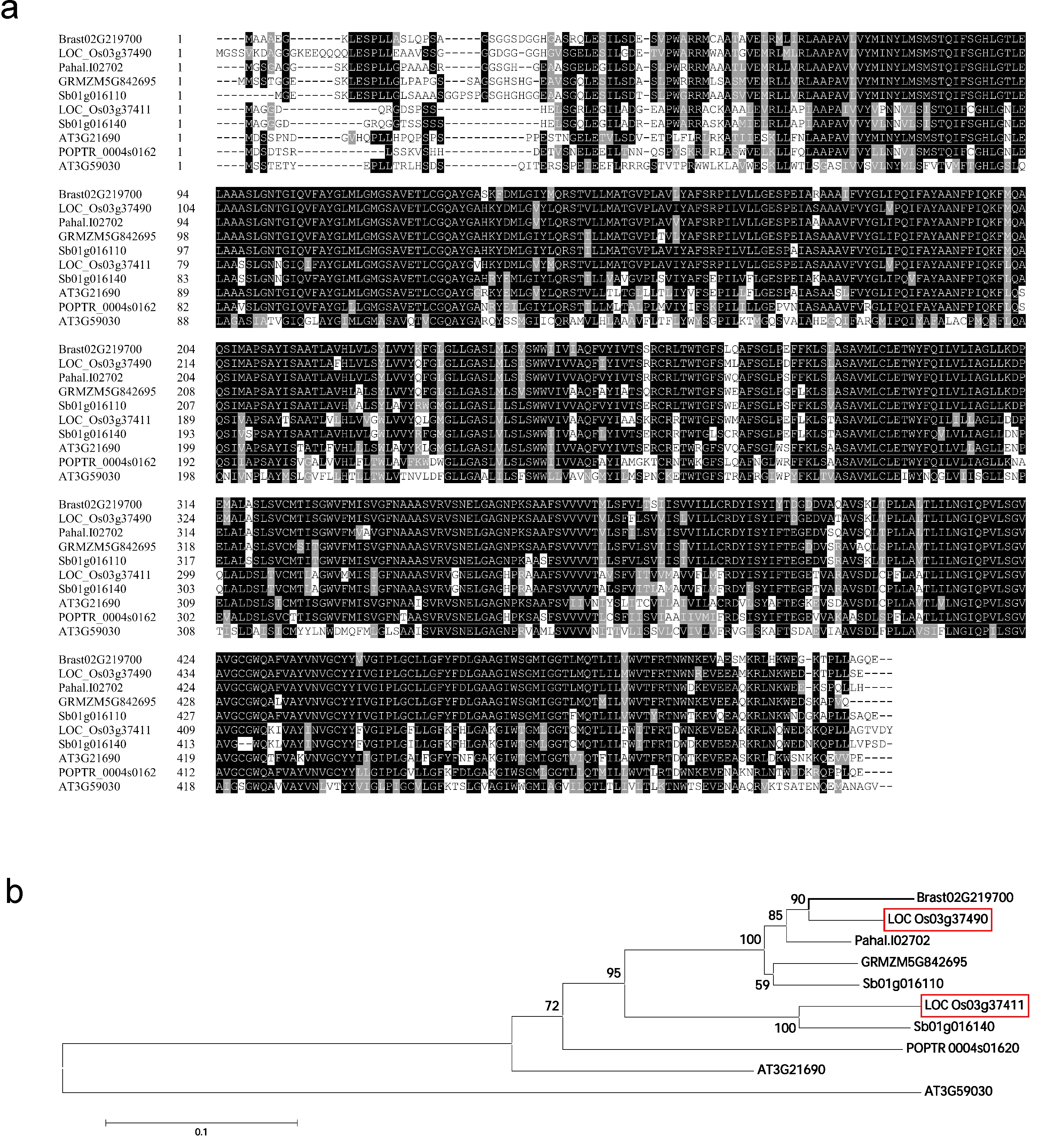

### Fig. S3.jpg

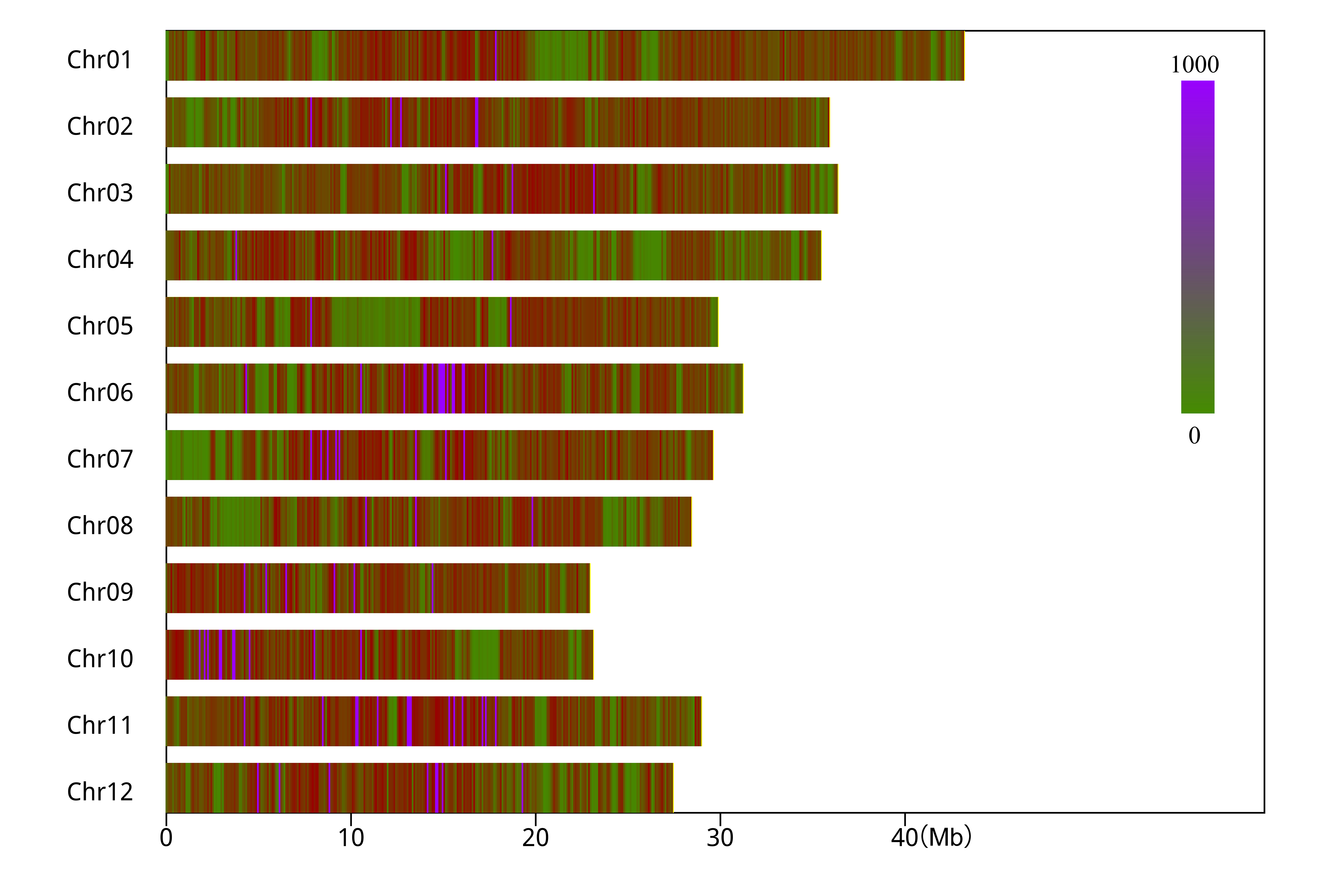

### Fig. S4.jpg

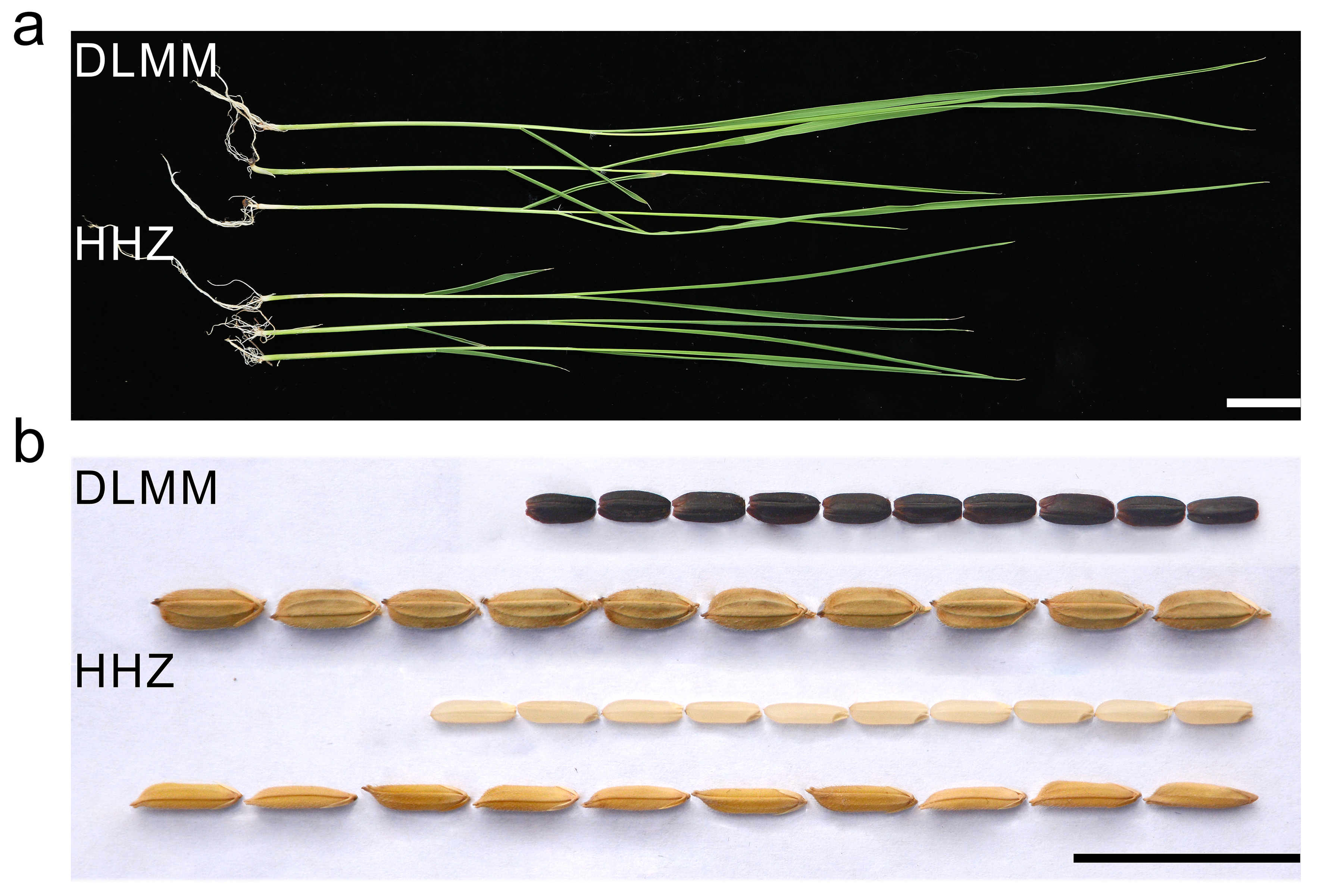

### Fig. S5.jpg

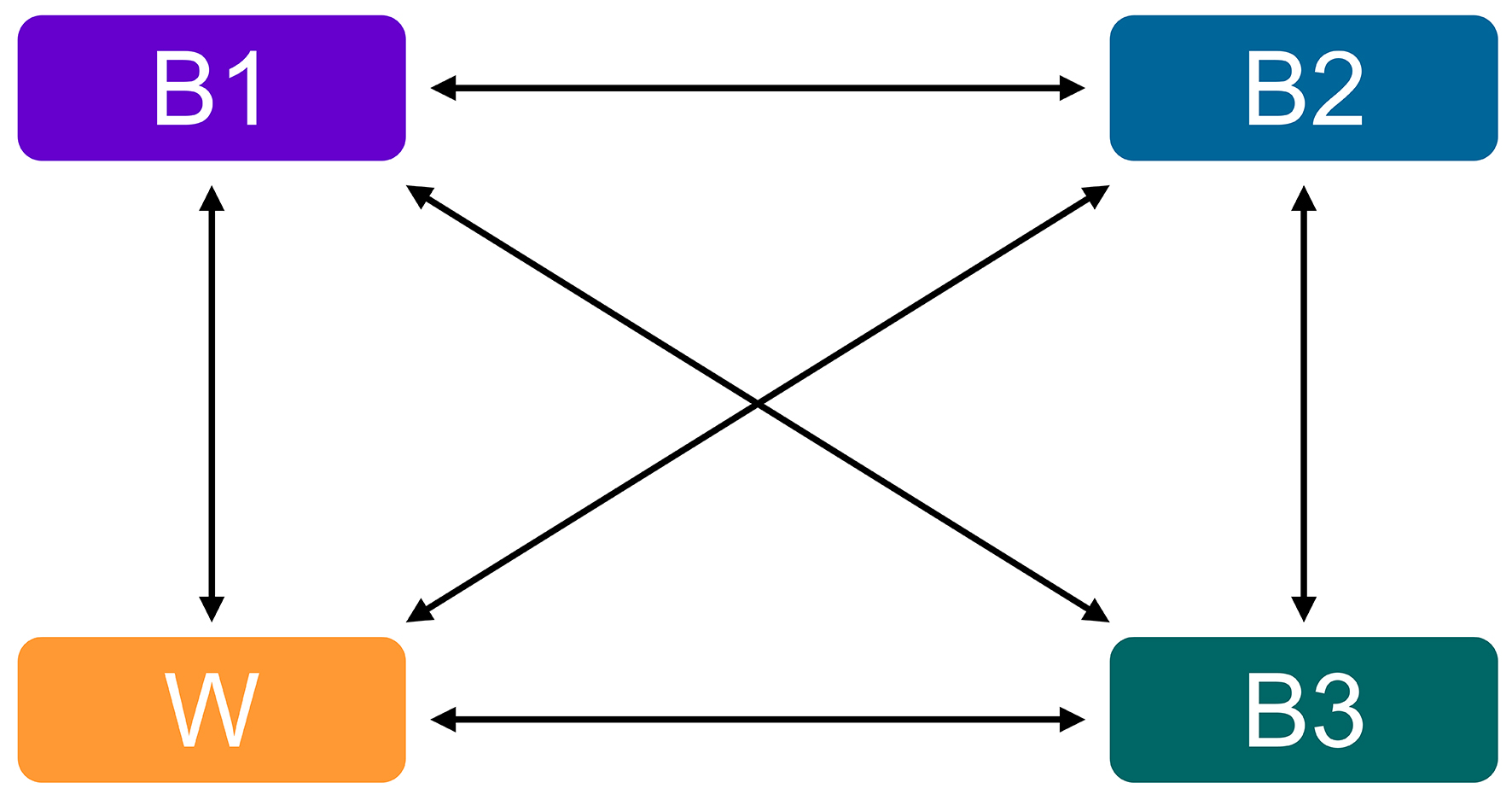

### Fig. S6.jpg

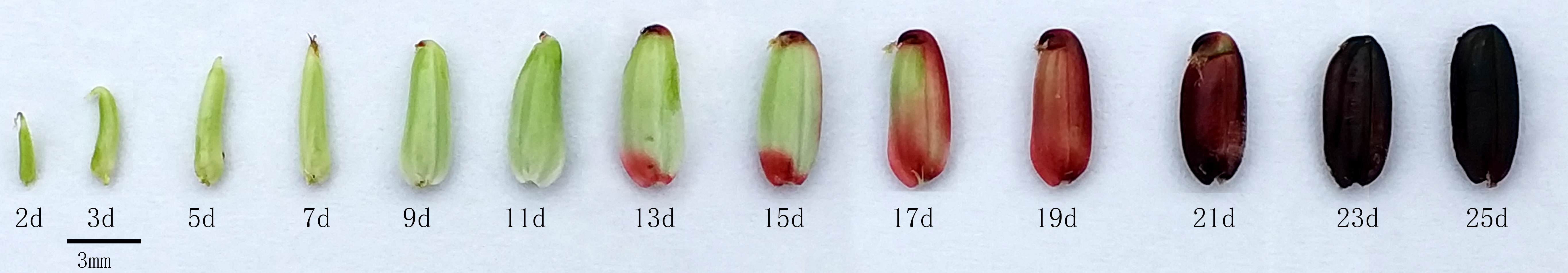
